# Apoptosis in the fetal testis eliminates developmentally defective germ cell clones

**DOI:** 10.1101/601013

**Authors:** Daniel H. Nguyen, Diana J. Laird

## Abstract

Many germ cells (GCs) are eliminated during development, long before differentiating to egg or sperm, but it is not clear why. Here, we examined how GC composition in the mouse fetal testis is altered by scheduled apoptosis during sex differentiation. Multicolored-lineage tracing revealed that apoptosis affects clonally-related GCs, suggesting that this fate decision occurs autonomously based on shared intrinsic properties. We identified extensive transcriptional heterogeneity among fetal GCs including an apoptosis-susceptible subpopulation delineated by high *Trp53* and deviant differentiation. Alternatively, the GC subpopulation most likely to survive was advanced in differentiation. These results indicate that GC developmental fate is based upon discrete and cell-heritable fitnesses and imply that a dichotomy between sex-differentiation and apoptosis coordinates the removal of developmentally incompetent cells to improve gamete quality. Evidence that GC subpopulations are in different epigenetic states suggests that errors in epigenetic reprogramming form the basis of aberrant differentiation and apoptotic selection.

**One sentence summary:** Germ cells undergo autonomous selection in the fetal testis to promote male differentiation

## INTRODUCTION

The transmission of genetic information in mammalia is tasked to the gametes - eggs and sperm – once organisms reach sexual maturation in adulthood. However, the formation and development of the gamete precursors, known as primordial germ cells (PGCs), begins far earlier during embryogenesis. PGCs are segregated from the soma early during fetal development and strictly maintained as a separate lineage (Ohinata et. al., 2009). To produce highly specialized gametes, PGCs first undergo a dynamic and complex development in the fetal period. In mice, this development entails a diverse range of cellular processes that begins with specification at E6.5 (Saitou and Yamaji, 2010) followed by a lengthy migration (Anderson et. al., 2000, Molyneaux et. al., 2001) to the site of the developing gonads. During this stage, PGCs interact with diverse niches that regulate behaviors like proliferation and motility (Cantú et. al., 2016). Concurrently, PGCs undergo extensive epigenetic reprogramming that includes histone modifications and DNA demethylation (Saitou et. al., 2012). After arriving in the gonads, PGCs expand and then sex differentiate into male and female germ cells in response to external signals (Menke et. al., 2003; Ohta et. al., 2012). Many elements of this tortuous process of establishing the germline are highly conserved (Barton et. al., 2016), reflecting the stringent requirements for creating cells able to carry out transgenerational inheritance.

The differential ability to correctly respond to developmental cues that guide these diverse processes, whether it be by directed migration or initiation of a differentiation, can create significant variation in the germ cell population. During migration, interactions with discrete niches produce vastly different PGC behaviors such as movement versus proliferation among PGCs at the same age (Cantú et. al., 2016). As a result, migratory outcomes can be highly heterogeneous depending on PGC position. Germ cell differentiation and survival are also influenced by numerous other variables including distribution in relation to signaling gradients (Doitsidou et. al., 2002) or heterogeneous expression of appropriate receptors (Morita-Fujimura et. al., 2009). Additionally, stochastic fluctuations in transcription can further alter a cell’s response to developmental challenges (Raj and Oudenaarden, 2008). Altogether, the complex interplay of these signals and responses can generate cellular heterogeneity and lead to divergent cell outcomes.

While the diversity of germ cell development can produce emergent heterogeneity, it also provides multiple opportunities for selection, such as the physical separation of PGCs depending on migratory competence (Cantú et. al., 2018). In the fetal period, apoptosis that occurs in both male and females further separates survivors from apoptotic losers, although the basis for this selection is unclear. Apoptosis has an important functional role: bypassing apoptosis results in male sterility (Knudson et. al., 1995) or increases the likelihood of reproductive defects in oocytes (Perez et. al., 1997). However, the severe requirement for apoptosis specifically in males suggests that programmed cell death is integrally linked with reproductive fitness (Aitken et. al., 2011). Prior to sex differentiation, apoptosis eliminates fetal germ cells that fail to migrate correctly (Stallock et. al., 2003). Fetal male germ cells also are subjected to an apoptotic wave between e13.5 to e17.5 (Coucouvanis et. al., 1993, Wang et. al., 1998). This apoptosis is dependent on the intrinsic apoptotic pathway involving the pro-apoptotic gene *Bax* but not the extrinsic *Fas*-dependent pathway (Runyan et. al., 2006). Importantly, male germ cell apoptosis is developmentally programmed and consistently occurs in the absence of typical cytotoxic insults such as radiation or chemical exposure. While previous studies of *Bax* mutant mice have investigated the effects of preventing apoptosis on adult spermatogenesis, little is known specifically about the fetal apoptotic wave and how it selects which germ cells survive or die.

Given their importance for genetic transmission, the identities of germ cells that survive to participate in gametogenesis have major reproductive implications for which genomes are propagated. While apoptosis eliminates germ cells from contributing to the germline, other developmental processes such as differentiation and proliferation may also positively select for certain germ cells. From specification through adult spermatogenesis, all these various developmental challenges have been shown in tetrachimeric mice to favor germ cells that originate from a certain founder population (Ueno et. al., 2009) and are largely responsible for the majority of spermatogenesis in the adult testis. Such results illustrate the evolutionary importance of developmental events in shaping the composition of the future germline.

Here, we investigate the specific wave of apoptosis during fetal germ cell development and determine that apoptosis is nonrandom; rather, it is clonally selective and eliminates germ cells with aberrant differentiation. We present a single-cell RNA-sequencing dataset during male differentiation that reveals heterogeneous differentiation and distinct subpopulations of germ cells with divergent fates for survival and apoptosis. We show that aberrantly differentiated germ cells are eliminated by apoptosis in a clonally-biased manner. In contrast, germ cells that are able to efficiently differentiate to a male state are resistant to apoptosis and characterized by activation of the machinery for piRNA biogenesis. We also distinguish these opposing germ cell states by their expression of epigenetically-regulated genes such as LINE-1, suggesting that differences in epigenetic reprogramming underlie clonal heterogeneity in survival and male identity. These results argue that apoptosis during male germline development functions as a quality control mechanism to promote efficient male differentiation and ultimately fitter gametes.

## RESULTS

### Germ Cell Apoptosis is Spatially Clustered in Fetal Testes

To detect the relative locations of dying germ cells during the wave of apoptosis, we employed a 3D imaging in situ approach on intact fetal testes. Wholemount immunostaining against a germ cell nuclear marker, TRA98, resolved individual germ cells. In conjunction with the late-stage apoptotic marker cleaved PARP (cPARP), we observed an acute increase in germ cell death from E12.5 through E14.5, consistent with previously published characterizations (Figure S1A) (18,19). Compared to other developing tissues throughout the embryo at similar timepoints (Foley and Bard, 2002) and germ cells in the fetal gonad at E11.5 before sex differentiation (Laird et. al., 2011), this represents more than a tenfold increase in apoptosis. In apoptosis-deficient *Bax* mice, we observed a 15% increase in overall germ cell number at E15.5, (data not shown) however this increase is likely limited by overcrowding of the germ cell niche and may underestimate the cumulative number of germ cells lost to apoptosis.

With the comprehensive spatial information provided by wholemount imaging, we noted that apoptotic germ cells displayed local clustering throughout the developmentally scheduled period of apoptosis (Figure 1A, Supplemental Movie 1). To confirm that apoptotic germ cells were indeed nonrandomly distributed, we extracted spatial coordinates for nuclei of apoptotic as well as all germ cells for three-dimensional analysis using Ripley’s K-function (Hansson et. al., 2013). From E12.5 through E14.5, apoptotic germ cells exhibited significantly increased spatial clustering compared to the distribution of all germ cells (Figure 1B). We did not observe a spatial bias at a tissue-wide level for apoptosis; rather, apoptotic clusters themselves were randomly distributed throughout the testis (Figure S1B), indicating that clustering was unrelated to tissue macrostructure or regional organization. Further investigation of the cellular environment immediately surrounding apoptotic germ cell clusters did not reveal any correlation with distance or ratio between germ cells and the supportive Sertoli cells (Mendis et. al., 2011) (Figure S1C-E) that have been thought to regulate germ cell survival in a paracrine manner. Since these extrinsic factors appeared to be unchanged between apoptotic and non-apoptotic environments, this suggested that the local environment does not strongly contribute to germ cell apoptosis. While we cannot completely exclude the influence of the local environment, these results indicated that intrinsic factors shared by germ cells in the observed clusters were instead primarily responsible for determining apoptotic likelihood.

**Figure 1.**
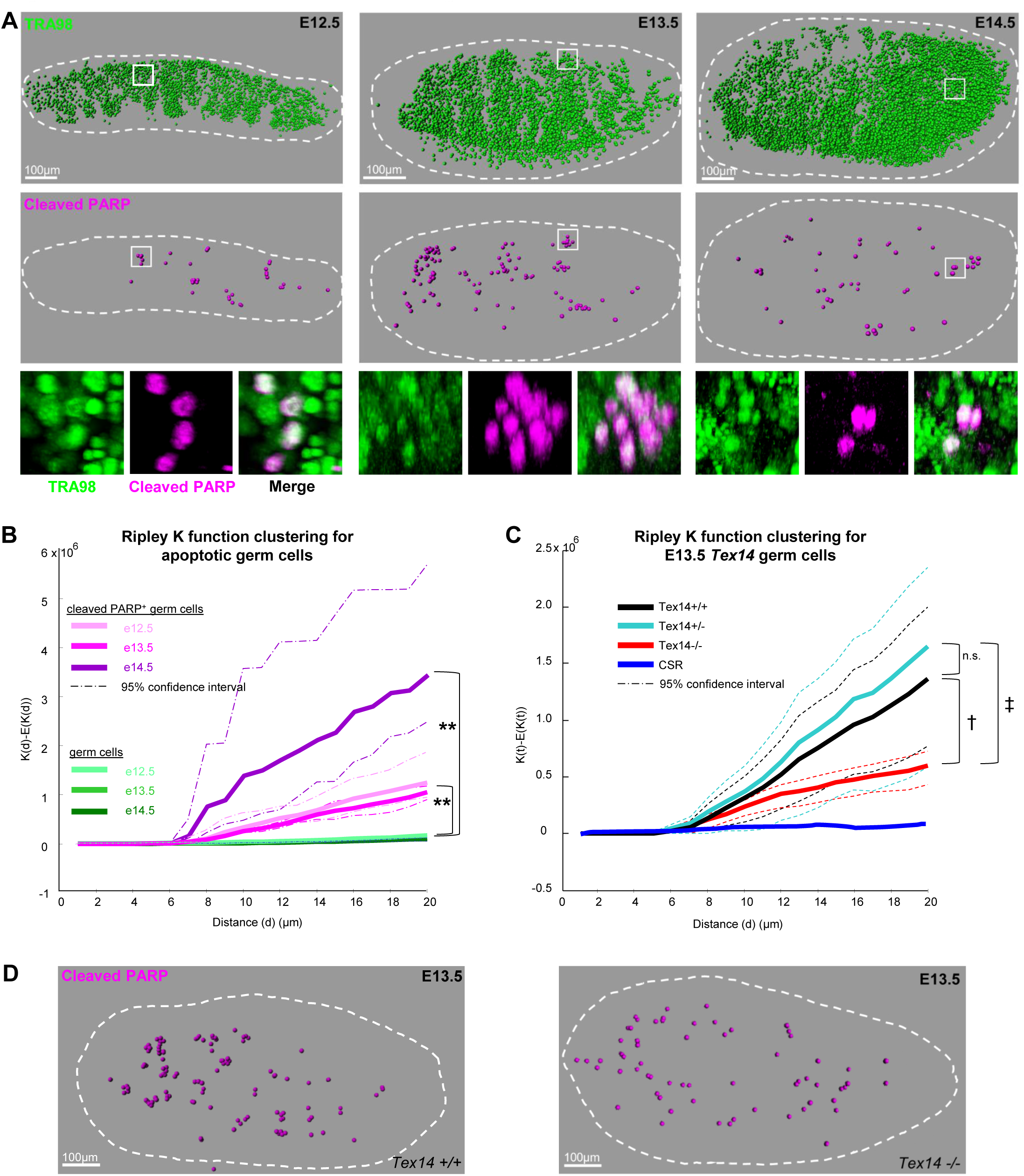
Germ cell apoptosis is spatially clustered in fetal testes. **A**, Wholemount imaging for germ cell nuclei and apoptosis enables spatial reconstructions of germ cell distributions in fetal testes. Germ cells from E12.5 through E14.5 testes were identified by nuclear marker TRA98 and individually mapped in three-dimensional space. Apoptosis within this germ cell population was detected by cleaved PARP. Bottom: Higher magnification examples of apoptotic germ cell clusters from wholemount imaging. B, Ripley K-factor analysis of spatial clustering of apoptotic germ cells (cleaved PARP^+^) from E12.5 – E14.5 testes compared to the extent of clustering of all germ cells (TRA98^+^), n=3 testes for each timepoint ** = p<0.05. **C,** Spatial clustering of apoptotic germ cells in *Tex14* compared to complete spatial randomness (CSR). n=2 testes for each genotype. **D,** apoptotic germ cell distributions in wholemount for *Tex14.* † p-value: 0.1716, ‡ p-value: 0.2476

A potential contribution to clustered apoptosis could come from phylogenetically conserved and specialized intercellular bridges that form between germ cells beginning at E10.5 due to incomplete cytokinesis (Lei and Spradling, 2013). The resulting germ cell cysts permit cytoplasmic sharing of components as large as organelles (Ventelä et. al., 2003) and could convey apoptotic signals to connected cells. To determine the contribution of these bridges to apoptotic clustering, we examined the distribution of apoptotic germ cells in *Tex14*^*-/-*^ mice that fail to form germ cell intercellular bridges (Greenbaum et. al., 2006). We found that the frequency of germ cell apoptosis was not significantly different from that of littermate wild-type fetal testis (Figure S1F). Importantly, clusters of apoptotic germ cells were still detected in mutant testis (Figure 1D). Though *Tex14* apoptotic clusters were smaller in cell number (Figure S1G), their continued existence indicates that local apoptotic potential is maintained in nearby disconnected cells. As cytoplasm is not shared among individual *Tex14*^*-/-*^ germ cells, diffusible apoptotic signals are therefore not necessary for the observed apoptotic clustering. This contrasts with clonal apoptosis in *Drosophila* spermatogonia that requires intracyst cytoplasmic sharing (Lu and Yamashita, 2017). It is likely that bridges in murine male germ cells coordinate some degree of timing and promote similar behaviors across interconnected cells of a cyst. However, nearby germ cells that descended from a common parent could continue to share apoptotic potential if it were an intrinsically heritable property that is present in all progeny cells regardless of interconnectivity.

### Multicolor Labeling Reveals Clonal Germ Cell Apoptosis

To investigate the lineage relationship among locally clustered apoptotic germ cells, we utilized two different multicolor reporter systems. By inducibly labeling multiple germ cell clones with distinct colors in the same testis, we are able to more comprehensively compare behaviors in neighboring, differently labeled clones than previous studies accomplished at single-clone density (Lei and Spradling, 2013). *Rosa26-Confetti* (Snippert et. al., 2010) or *Rosa26-Rainbow* (Rinkevich et. al., 2011) mice were crossed to *Pou5f1-CreERT2* (Greder et. al., 2012) mice and given a single dose of Tamoxifen at E10.5 to induce recombination in germ cells at approximately E11.0 after Tamoxifen metabolism. E10.5 was chosen as an ideal timepoint because it represents the end of migration when germ cells begin proliferation in a sessile state (Figure 2A). By E13.5, germ cell clones comprising one of 3 or 4 different colors grew to a mean size of 8 (Figure Supplemental Movie 2). Clone sizes were consistent with previously described cyst formation (Lei and Spradling, 2013) and clones were connected by TEX14-positive structures, confirming their cytoplasmic sharing (Figure Supplemental movie). From E12.5 through E14.5, we observed that individual clusters of apoptotic germ cells were the same color (Figure 2B) and that apoptotic cells lay within the boundary occupied by a single clone (Figure S2). We categorized the distribution of apoptosis in clonally labeled local clusters and found that apoptosis was strictly monoclonal (Figure 2C), even when differently labelled clones were adjacent and interspersed. The absence of polyclonal apoptosis in a local cluster strongly argues against an extrinsic basis for apoptosis and instead suggests that intrinsic, clonally-shared properties could predispose subpopulations of germ cells toward either survival or apoptosis.

**Figure 2:**
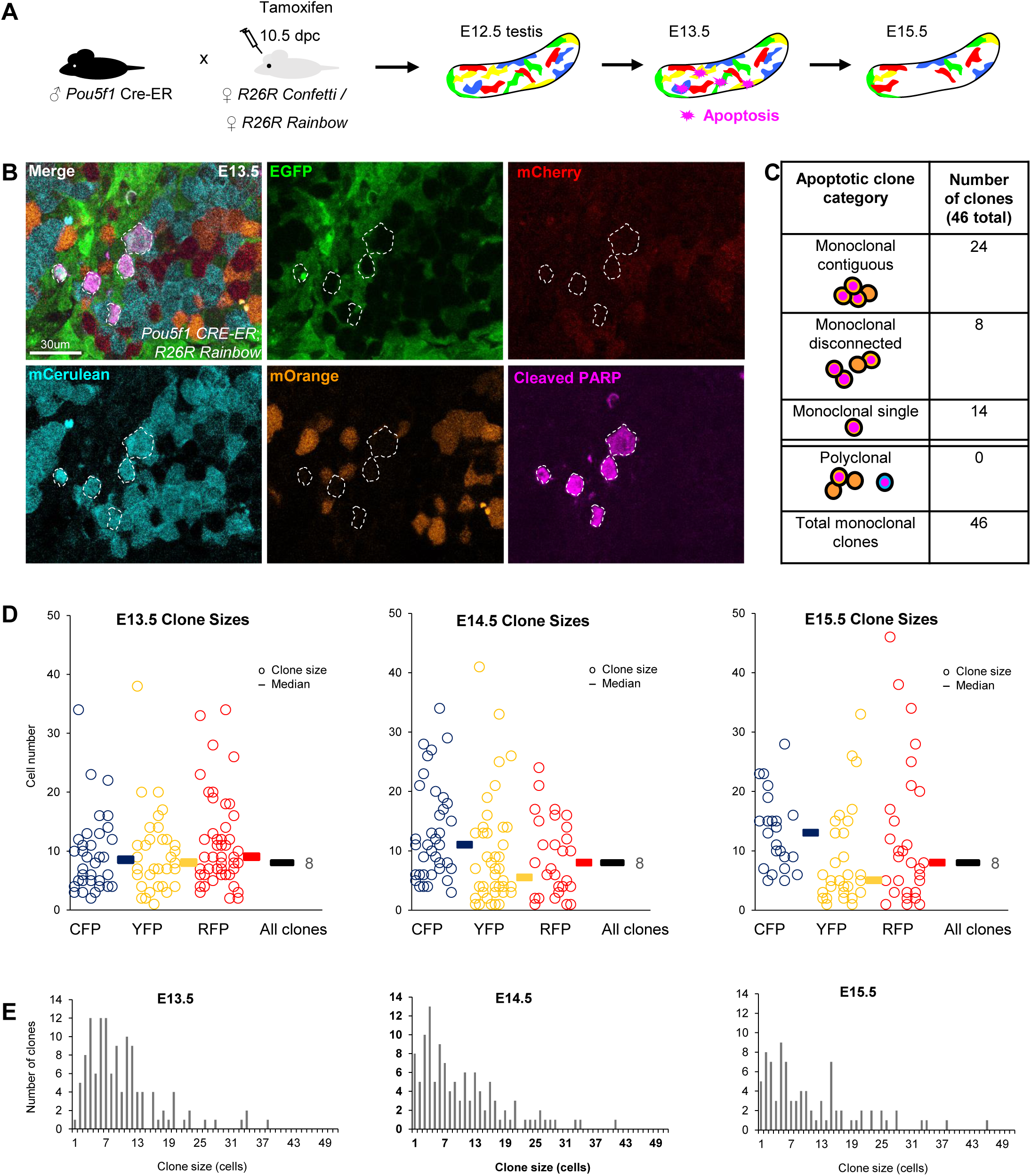
Multicolor clonal labeling reveals clonal apoptosis during the fetal apoptotic wave. **A,** Experimental scheme for specific labeling of germ cells with multicolor reporters *Confetti* or *Rainbow*. **B**, Apoptotic cluster in E13.5 *Rainbow* testes containing similarly labeled (mCerulean) germ cells. **C,** Clustered *Confetti* or *Rainbow* apoptotic germ cells categorized by cluster organization and clonality. **D,** Clone size measurements during apoptotic wave. Each clone is represented by a circle with the corresponding color. Median clone sizes are shown for each color as well as for all colors combined (black bars, median clone size value adjacent). **E,** Histogram displaying clone size distributions throughout apoptotic wave. One-way ANOVA for all 3 stages p-value =0.806248.

We observed instances where an entire clone was cPARP positive as well as partially cPARP positive; however, this detection of apoptosis is limited by the transience of cPARP in dying cells and the rapid subsequent cellular breakdown. To evaluate whether apoptosis cumulatively affects all cells in clones or only a subset, we compared the size of clones across development: at E13.5 following mitotic arrest of germ cells (Western et. al., 2008) and at E15.5 concluding the apoptotic wave. A simple model of clone size dynamics during apoptosis predicts that partial death of a clone would manifest as a decline in average size of clones. However, we observed a consistent mean clone size of 8 from E13.5 until E15.5 past the conclusion of the apoptotic wave in our system (Figure 2D, 2E). This result concurs with the size of clones reported in experiments where only one single clone is labeled per testis (Lei and Spradling, 2013), demonstrating the scalability of a multicolor comparative approach. Together, constant clone size and confined clonal apoptosis argue that germ cells in the fetal testis are eliminated based upon mitotically-heritable properties. Furthermore, the distinct clonal outcomes in regard to survival suggest that the germ cell population contains significantly heterogeneous subpopulations.

### Single-cell RNA Sequencing Identifies an Apoptosis-Poised Germ Cell Subpopulation

To identify the molecular basis of the observed heterogeneity distinguishing apoptotic from non-apoptotic germ cells, we performed single-cell RNA-sequencing (scRNA-seq) of wild-type germ cells from E13.5 testes at the peak of apoptosis. Single-cell transcriptomics of purified germ cells from a single timepoint should reveal heterogeneity of cell states within the germ cell population and allow for the detection of an apoptotic state that would otherwise be masked in a bulk analysis (Patel et. al., 2014). As multicolored germ cell lineage labeling studies established that apoptosis is a clonal behavior, we reasoned that the expression of pro-apoptotic genes would be detectable as a discrete cluster in a single-cell analysis.

We purified *Pou5f1-Oct4*GFP+ germ cells from testes at E13.5 for scRNA-seq. 2,585 germ cells clustered into 9 distinct subpopulations based on their global transcriptional profiles (Figure 3A). We identified markers that characterized each subpopulation (Supplemental Table 1) and examined them for expression of apoptosis-related genes. The majority of germ cells at E13.5 maintained high levels of the pro-apoptotic gene *Bax*, consistent with prior observations (Runyan et. al., 2006) (Figure 3B, Figure S3A). Subpopulation 6 was notable for expressing the highest levels of *Trp53* among all other subpopulations. Elevated expression of *Trp53* and its protein product P53 is known to disadvantage cells in competitive transplant assays in blood as well during embryogenesis (Bondar and Medzhitov, 2010; Bowling et. al., 2018; Zhang et. al., 2017). Downstream targets of *Trp53* were confirmed to be increased in subpopulation 6 (Figure S3B). As p53 plays an established role in apoptosis (Haupt et. al., 2003), we examined the expression of other pro-apoptotic genes across the 9 subpopulations (Figure 3B). Subpopulation 6 cells expressed the highest levels of transcripts encoding genes involved in germ cell apoptosis such as *Bax* and *Bad* (Figure 3B) (de Felici et. al., 1999). The expression profile of subpopulation 6 in regard to pro-apoptosis genes is characteristic of an apoptotic state; we henceforth refer to population 6 as apoptosis-poised (AP) germ cells. We noted that AP-germ cells represented 7% of the germ cell population assessed in our scRNA-seq dataset, which is consistent with the observed percentage of apoptotic germ cells from wholemount analysis (Figure S3B).

**Figure 3:**
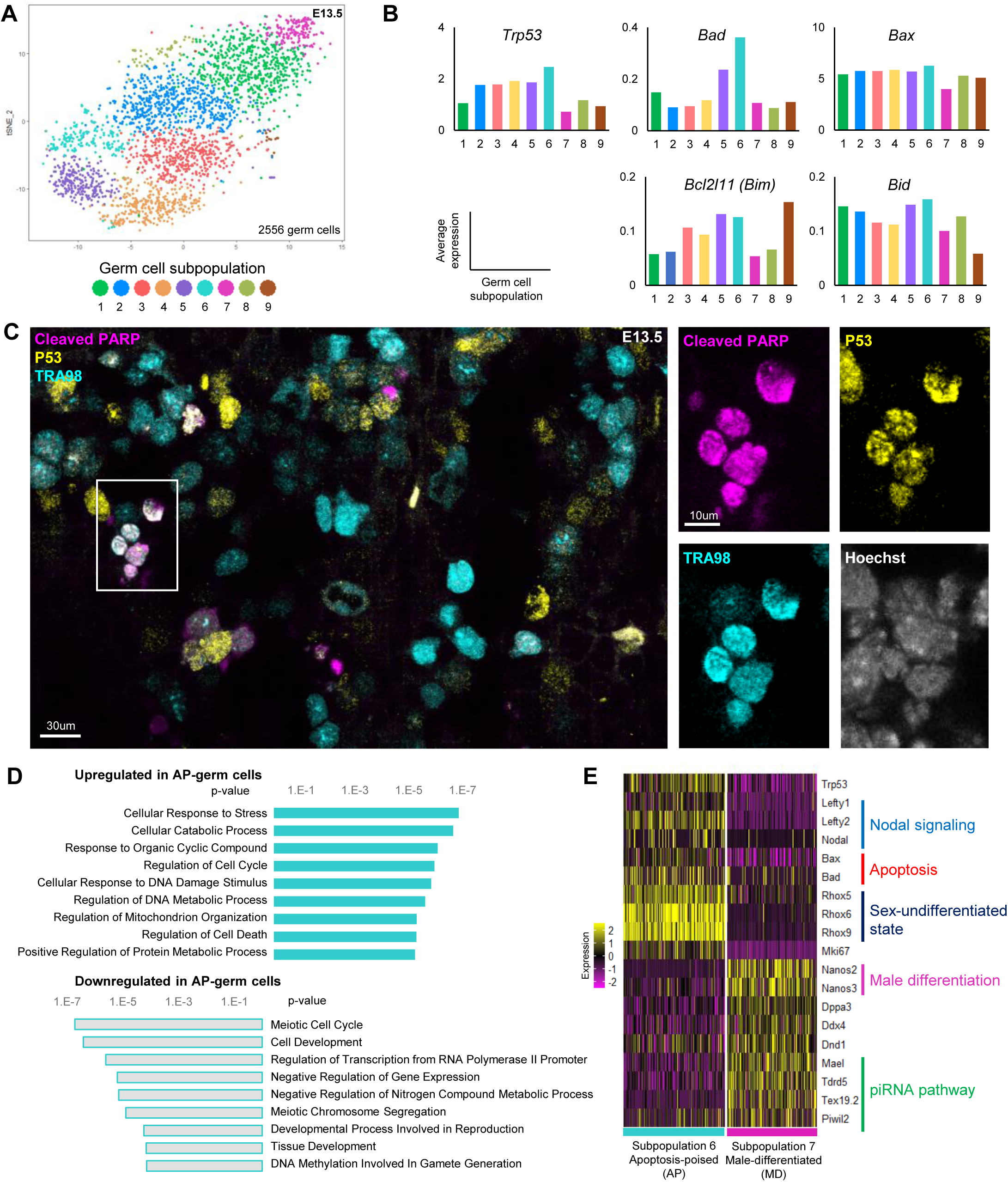
Single cell RNA sequencing of male germ cells identifies an apoptotically-poised subpopulation. **A,** Transcriptional heterogeneity and clustering of 2,556 E13.5 germ cells by tSNE plot organizes cells into 9 populations. **B,** Expression of apoptosis-related genes by individual cells of each of the nine germ cell states. **C,** Immunofluorescence staining for coexpression of p53 and apoptosis (cleaved-PARP) in E13.5 testis sections. **D,** Gene ontology of transcriptional markers upregulated or downregulated in the *Trp53-*high germ cell state 6 compared to all other clustered populations. **E,** Expression heatmap for genes related to germ cell death and differentiation in apoptosis-poised (AP) germ cells (subpopulation 6) and apoptosis-resistant male-differentiating (MD) germ cells (subpopulation 7).

Given that *Trp53* was identified as a significant marker of AP-GCs, we examined P53 protein expression in E13.5 germ cells during the peak of apoptosis. Immunostaining revealed levels of P53 to be heterogeneous among germ cells and elevated in germ cells undergoing clustered apoptosis (Figure 3C). The clustered pattern of p53 heterogeneity is consistent with clonal expression.

### Apoptosis-Poised Germ Cells Exhibit Aberrant Male Differentiation

Gene ontology (GO) analysis of the other upregulated genes specifically distinguishing AP-germ cells (TABLE S1) found enrichment for cell death and stress pathways (Figure 3D). Conversely, genes that were specifically downregulated in AP-germ cells represented pathways involved in germline differentiation. Altogether, GO analysis suggests that apoptosis and germline differentiation represent opposing germ cell states at E13.5. To confirm this, we investigated if a subpopulation contrasted with AP-germ cells for expression of apoptosis genes. Although most germ cell subpopulations at E13.5 exhibited high levels of pro-apoptosis transcripts, we noted that a single subpopulation – subpopulation 7 – expressed the lowest levels of apoptosis-associated markers such as *Trp53, Bax* and *Bad* (Figure 3B) compared to all other subpopulations. The reciprocal expression of apoptosis genes between AP-germ cells and germ cells in subpopulation 7 positions these two groups of germ cells as potentially opposite states; therefore, we sought to contrast these subpopulations to characterize deficiencies in the AP-germ cells.

In addition to apoptosis, other transcriptional differences between AP-germ cells and subpopulation 7 germ cells provided further insight into their divergent states. AP-germ cells were uniquely defined by high levels of *Rhox*-family genes (Figure 3E). Though ordinarily absent in E13.5 male-differentiated germ cells, *Rhox* genes are expressed in both male and female germ cells prior to E13.5 and decrease sharply after E12.5 in males (Maclean et. al., 2005) in association with male differentiation. The persistence of *Rhox* expression in the E13.5 AP-germ cells could therefore indicate a developmentally delayed state.

This hypothesis was bolstered by the high expression of Nodal signaling-responsive components including *Lefty2* in the AP-germ cell subpopulation (Fig 3E). Nodal signaling is normally active in E12.5 male germ cells that have not fully committed to the male lineage but its duration is transient and it must be extinguished in order for male differentiation to proceed efficiently; negative feedback quickly shuts Nodal signaling down by E13.5 (Spiller et. al., 2012). Distinctively high *Lefty2* expression in AP-germ cells relative to all other E13.5 subpopulations therefore reflects atypically persistent Nodal signaling and disrupted male differentiation in these cells.

Consistent with defective differentiation, transcripts that should be expressed in advanced male GCs, including *Nanos2* (Saba et. al., 2014) and piRNA-regulating genes (Molaro et. al., 2014), were instead downregulated in AP-germ cells (Fig 3E). In contrast, subpopulation 7 was defined by the highest expression of *Nanos2* and PIWI-piRNA genes that are characteristic of an advanced male-differentiated (MD) germ cell state.

Given that AP-germ cells are characterized by high expression of apoptosis-related genes and an absence of male-differentiation genes, we examined the relationship between markers of apoptosis and differentiation at a population-wide level. The expression of *Trp53,* a marker for AP-germ cells, and male-differentiation gene *Nanos2* was inversely distributed in the entire E13.5 single-cell RNA-seq dataset (Figure 4A). Correspondingly, we observed a reciprocal relationship between *Trp53* and *Nanos2* transcript levels in male germ cells at E13.5 by RNA in-situ hybridization (Figure 4B). Further supporting the dichotomous relationship between apoptosis and differentiation is the reciprocal expression of *Nanos3* between MD and AP subpopulations; this suppressor of germ cell apoptosis (Suzuki and Saga, 2008) is highest in MD germ cells (Fig S3C).

**Figure 4:**
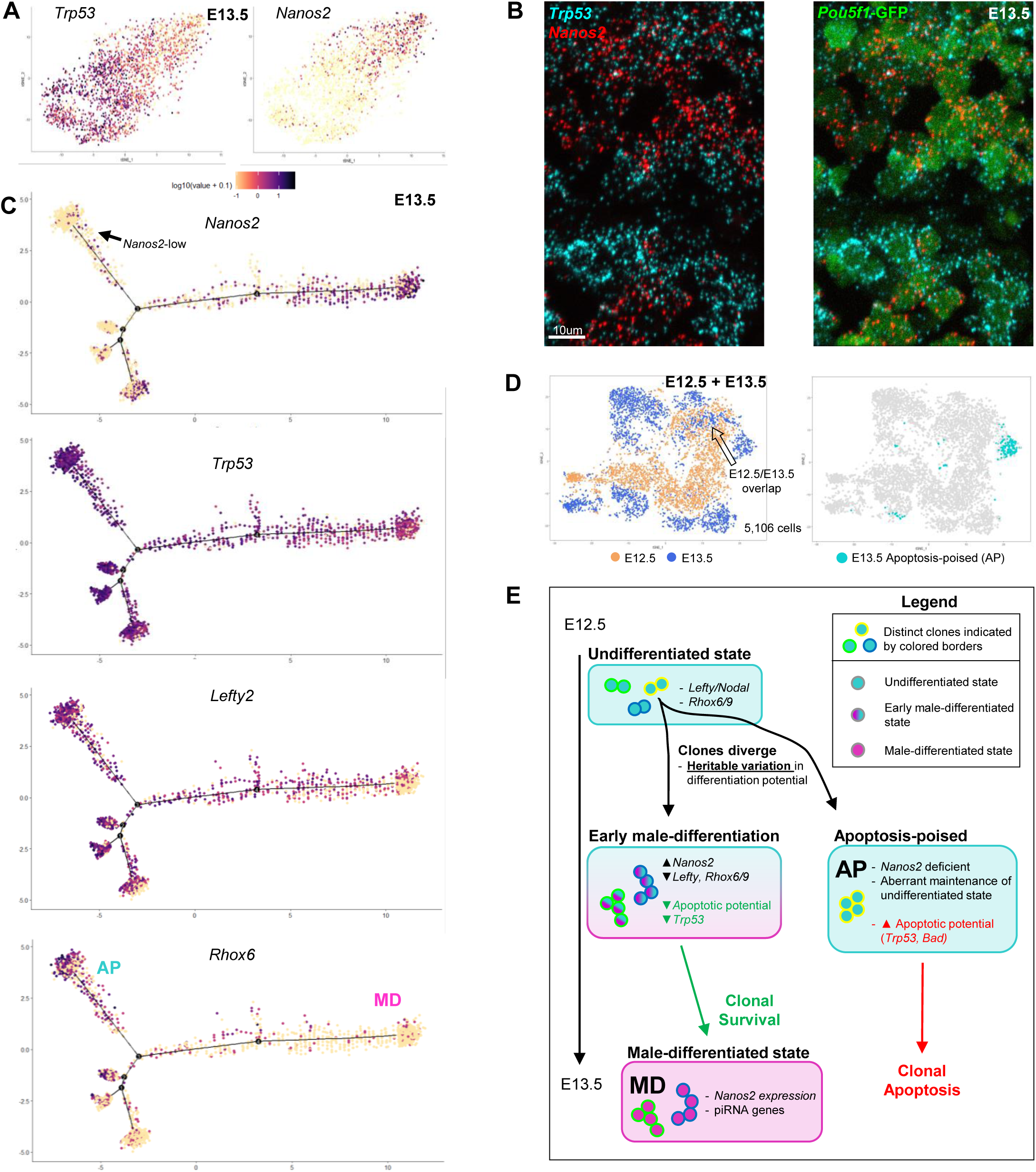
AP germ cells exhibit aberrant male sex differentiation at E13.5. **A,** *Trp53* expression in E13.5 single cells compared to differentiation marker *Nanos2*. B, Reciprocal expression of *Trp53* and Nanos2 mRNA in E13.5 cells by RNAscope. **C,** Psuedotime distribution of E13.5 single cells with expression for markers of male-differentiating (MD) germ cells (*Nanos2*) versus markers of an undifferentiated state (*Rhox6*) or apoptotic state (*Trp53, Bad*) that characterizes the apoptosis-poised population (AP). Arrow: Distinct *Nanos2-*low germ cell subpopulation with similar transcriptional profile to AP-germ cells. **D,** tSNE plot of combined E12.5 and E13.5 populations with age identified by color. Arrow: Overlapping E13.5 subset with E12.5 germ cells. Left: (Apoptosis-poised (AP)) separately identified on the combined E12.5-eE3.5 tSNE distribution. **E,** Model for coordination of clonal apoptosis with male differentiation. Clonal variation in differentiation potential results in divergent clonal fates: germ cells normally activate male differentiation programs marked by *Nanos2* but a subset of clones aberrantly maintain an undifferentiated state characterized by expression of pro-apoptotic genes that ultimately results in clonal death.

### AP-Germ Cells Deviate From a Pseudotime Trajectory of Normal Male Differentiation

We used pseudotime analysis to identify developmental trajectories among E13.5 GCs and place AP-germ cells in a differentiation context. The inferred trajectory contains *Nanos2-*positive cells along a main axis with several branches, with that marked by an arrow notable for an absence of *Nanos2* (Fig. 4C). This terminal branch also exhibits uniquely high *Rhox6/9*, as well as relatively elevated *Trp53* and *Lefty2* compared to other germ cells. Based on the expression of these markers, the resemblance of this branch to the AP-germ cell subpopulation suggests that the AP-germ cell state is a deviation from normal male differentiation.

The other germ cells that populate the pseudotime trajectory express *Nanos2*, with the highest *Nanos2*-expressing cells anchored one end of this main axis, suggesting that these are the most advanced male-differentiated germ cells. These cells are also distinguished by an absence of *Lefty2*, confirming their identity as MD-germ cells. The remaining cells on the main axis are *Nanos2-*positive but also *Lefty2*-positive, suggesting that they maintain some degree of early undifferentiated germ cell character and are likely an intermediate subpopulation transitioning between E12.5 to E13.5. Importantly, this analysis raises the possibility that the AP-germ cell state is a deviation from normal male differentiation rather than an intermediate state.

### AP-Germ Cells Are Transcriptionally Distinct from E12.5 Germ Cells

The observed heterogeneity in *Nanos2* across the entire germ cell population at E13.5 suggested that subpopulations vary in the extent of male-differentiation, with MD-germ cells representing an advanced extreme. In contrast, *Nanos2*-low germ cells could represent an earlier state of differentiation more similar to E12.5 germ cells. We profiled single-cell transcriptomes of male germ cells at E12.5 in order to directly interrogate the transcriptional overlap between E13.5 AP-germ cells and E12.5 germ cells. We computationally pooled both E12.5 and E13.5 datasets and batch-corrected by timepoint identity. Transcriptional clustering on the merged datasets revealed that the majority of germ cells at E12.5 and E13.5 cells were transcriptionally distinct in accordance with their embryonic stage (Figure 4D). This was confirmed by the stage-specific expression of germ cell developmental markers in each population (Fig S4). Maturation markers such as *Nanos2* were entirely absent in germ cells at E12.5. By contrast, markers of a sex-undifferentiated state – most notably *Trp53* and pro-apoptotic transcripts – were more uniformly high among all E12.5 cells. A subset of E13.5 germ cells overlapped with a subset at E12.5 and could represent an intermediate state with slightly delayed differentiation. However, AP-germ cells were not associated with this overlapping population and instead remained transcriptionally distinct from all other E12.5 cells. This difference suggests that the AP-germ cells are neither sex-differentiated E13.5 cells nor undifferentiated E12.5 cells, but instead a deviation from the normal developmental trajectory that ultimately terminates in apoptosis (Figure 4E model). The clonal nature of AP-germ cell apoptosis implies that aberrant differentiation potential arose from a cell-heritable defect that enforces a separate apoptotic fate for AP cells.

### Aberrantly Differentiated Germ Cells Are Retained in the Absence of Apoptosis

The expression of developmentally-inappropriate genes in the subpopulation of germ cells most likely to die suggests that the wave of scheduled apoptosis eliminates germ cells that fail to differentiate properly. A prediction of this hypothesis is that mis-differentiated germ cells bearing the hallmarks of the AP subset would persist in the absence of apoptosis. Germ cells with abnormal morphology that arrest prior to meiosis were observed in mice with inactivation of *Bax* or overexpression of *Bcl-x*, although little is known about the consequence of *Bax* elimination in the fetal period (Rucker et. al., 2000). To evaluate the germ cell identities that persist when apoptosis is prevented, we examined testes from *Bax*^*-/-*^ mice at E15.5, following the wave of fetal apoptosis when germ cells in wild-type tissue are destined for apoptosis would be removed.

Based on the distinctive expression of *Lefty2* in AP-germ cells at E13.5, we compared the dynamics with a LEFTY1/2 antibody. Whereas nearly all wild-type germ cells were LEFTY+ at E12.5, this frequency decreased sharply at E13.5 and continued to decrease at E14.5 (Figure 5A). By contrast, in *Bax*^*-/-*^ testes at E15.5, the percentage of Lefty-positive germ cells was over ten-fold higher compared to wild-type and heterozygous littermates (Figure 5B). The declining levels of LEFTY-positive germ cells during the wave of apoptosis in wild-type, together with the incomplete loss in the absence of apoptosis is consistent with the idea that successful male differentiation extinguishes Nodal signaling whereas failed differentiation is accompanied by persistent Nodal signaling. The LEFTY-expressing germ cells that are retained in *Bax*^*-/-*^ at E15.5 are distributed in a clustered manner typical of clonal expression (Figure 5c). Given that germ cell apoptosis is clonal during this period, this observed pattern in *Bax*^*-/-*^ is suggestive of clonal survival of AP-germ cells through E15.5.

**Figure 5:**
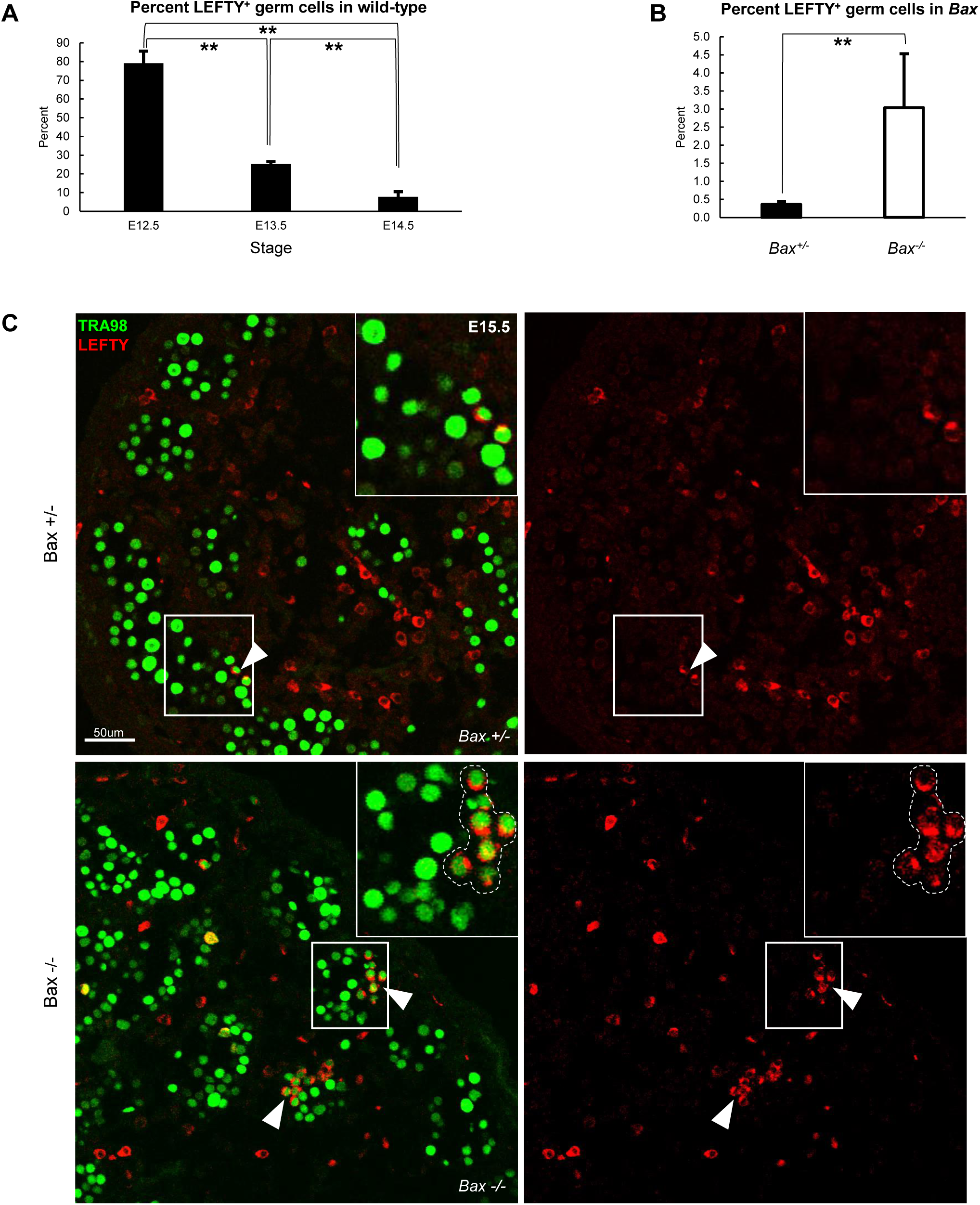
Aberrantly differentiated germ cells are retained in the absence of apoptosis. **A,** Percent Lefty expression in germ cells during male differentiation from E12.5-E14.5. ** = p<0.05, one way ANOVA. **B,** Percent LEFTY expression in Bax heterozygous versus homozyogous KO germ cells at E15.5. N=2 for each genotype. ** = p<0.05., Student’s t-test. **C,** Immunofluorescent detection of LEFTY expression in germ cells from *Bax* heterozygous versus homozygous KO testes at E15.5. White arrowheads highlight LEFTY-positive germ cell groups.

### Epigenetically-regulated Genes Underlie Germ Cell Heterogeneity

The dichotomous AP-germ cell and MD-germ cell states with divergent cell identities and fates highlights the heterogeneity among germ cells at E13.5. Furthermore, the clustered manner of AP-germ cell-associated *Lefty* expression combined with the clonality of apoptosis argue that apoptosis-related heterogeneity results from cell-heritable differences among germ cell subpopulations. Although spontaneous genetic mutations are a potential source of de novo heritable heterogeneity, the mutation rate in mice is too low to account for the percentage of apoptosis measured in germ cells. (Milholland et. al., 2017). However, cell-heritable changes can also arise from epimutations – de novo differential epigenetic marks such as variable methylation – that are transmitted to daughter cells through proliferation. The genome-wide DNA demethylation that occurs between E7.5 through E12.5 during germ cell development provides a significant opportunity for epimutations to arise and produce heritable heterogeneity among germ cell clones.

As suggested by the magnitude of epigenetic reprogramming in this period, the precise extent of demethylation, particularly at specific loci, is highly influential in controlling germ cell fate. Deletion of DNA methyltransferase *Dnmt1* leads to precocious demethylation and inappropriate male differentiation (Hargan-Calvopina et. al., 2016). Loss of *Tet1*, a 5mC oxygenase involved in DNA demethylation, also disrupts expression of genes important for germ cell sex differentiation (Hill et. al., 2018). Recent studies identified a set of 45 genes involved in germ cell differentiation that are tightly controlled by methylation at their promoters (Hill et. al., 2018). The stage-appropriate expression of these germline reprogramming responsive (GRR) genes can therefore indicate the degree of DNA demethylation of the cell.

In comparing the levels of GRR gene transcripts between the two E13.5 germ cell states we identified as diametrically male-differentiated, we found that a significant number of GRR genes overlapped with markers of the MD-germ cell state (Figure 6A). This overlap confirms the male-differentiated identity of MD-germ cells and suggests that the epigenetic state of MD-germ cells is permissive for male differentiation. Conversely, AP-germ cells were characterized by the depletion of GRR expression, with several GRRs identified as uniquely downregulated markers of the AP-germ cell subpopulation. This deficient GRR expression indicates that epigenetic reprogramming is incomplete in AP-germ cells and raises the possibility that germ cell heterogeneity is due to variation in epigenetic states.

**Figure 6:**
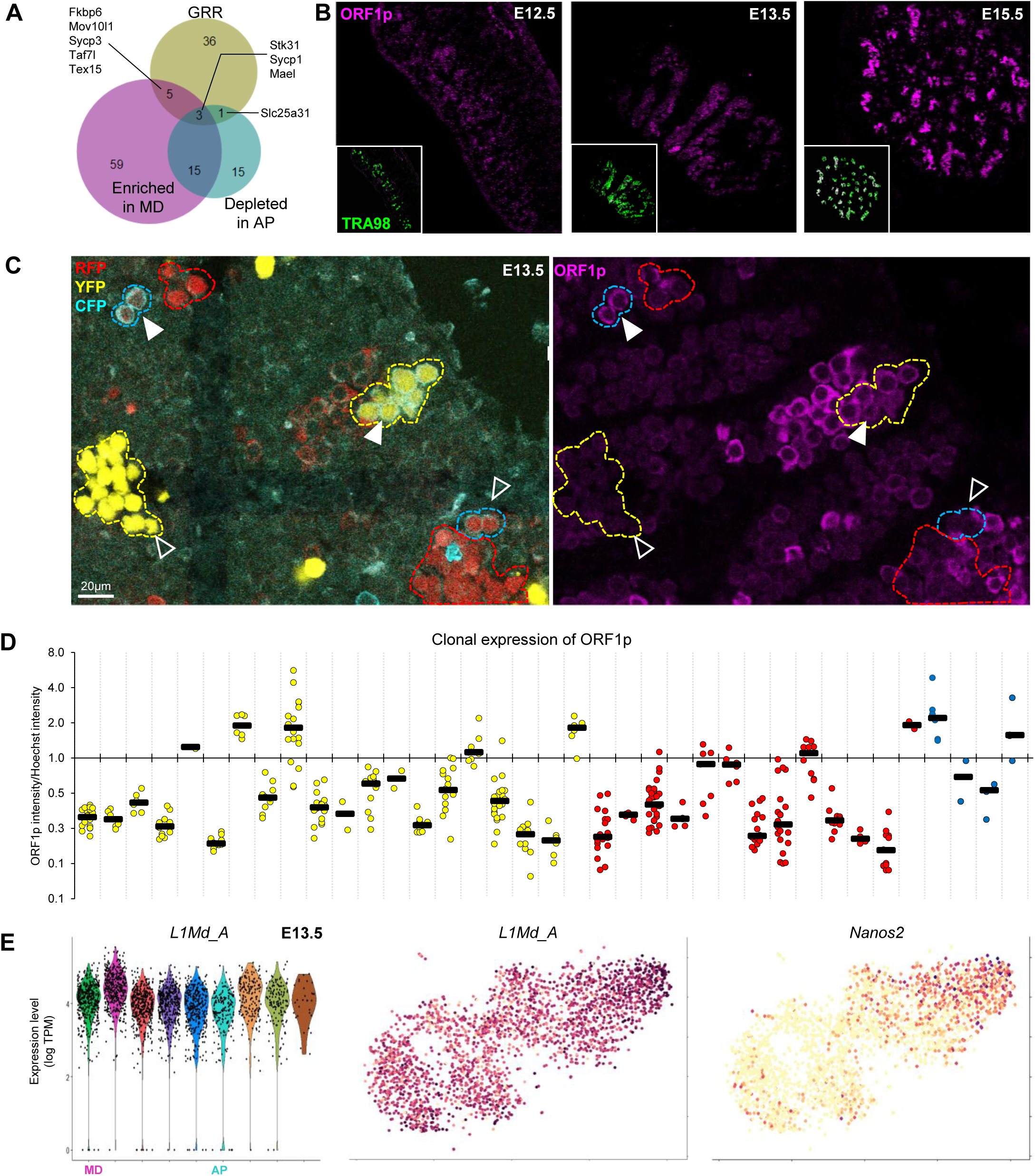
Epigenetically-regulated genes underlie germ cell heterogeneity. **A,** Overlap of total of 45 germline reprogramming-responsive genes (GRR) in MD population as well as their absence in the AP population. p-values GRR & AP-germ cells: **5.314e-07**, GRR & MD-germ cells: **3.120e-12**, AP-germ cells & MD-germ cells: **1.813e-36**. **B,** LINE1 expression dynamics during male differentiation detected by ORF1p antibody. **C,** ORF1p heterogeneity is clonally based as detected by *Confetti* labeling. **D**, Quantification of ORF1p immunofluorescence in individual cells of clones compared to intraclonal levels. **E,** Single cell repeat element RNA-seq identifies prominent LINE1 L1_MdA as a marker of the Nanos2-high MD-germ cell subpopulation

### TE Expression Is Heterogeneous Between Clones

In addition to activating GRR expression to initiate male differentiation, epigenetic reprogramming in fetal germ cell development is also known to regulate the expression of transposable elements (TE). TEs are repetitive elements that are uniquely regulated in the germline by epigenetic means as well as the piRNA pathway. Methylation at TEs is a major repressive strategy and TEs have been noted to remain relatively more methylated even during the massive global demethylation that occurs during fetal germ cell development (Seisenberger et. al, 2012). Since demethylation can render the host genome vulnerable to activation of TEs, expression of piRNA pathway components is coordinated with this demethylation. Given the discrete differentiation states we were able to identify in E13.5 male germ cells, we sought to investigate how TE expression may vary among these subpopulations, particularly because observed piRNA-related genes such as *Mael, Stk31*, and *Mili*, are differentially expressed in AP-germ cells versus MD-germ cells.

We examined TE expression in fetal germ cells by detecting the LINE-1 protein product ORF1p as germ cells progressed from a male-undifferentiated to differentiated state (E12.5-E15.5). LINE-1s are the predominant active TE expressed in germ cells and their regulation by piRNAs is well established (Aravin et. al., 2008). ORF1p expression progressively increased as germ cells complete male differentiation (Figure 6B). While ORF1p is entirely absent prior to the initiation of male differentiation at E12.5, small clusters of ORF1p-positive germ cells are detectable at E13.5. By E15.5 when male differentiation is complete, nearly all germ cells are ORF1p-positive. The clustered ORF1p expression is reminiscent of the clonal germ cell apoptosis also observed at these timepoints. To ascertain if ORF1p expression is similarly clonal, we measured ORF1p levels on a clonally labeled background at E13.5 when these heterogeneous ORF1p clusters are first observed (Figure 6C). Within each clone, constituent cells expressed similar levels of ORF1p, which implies that the ORF1p expressing state is stably heritable among related germ cells (Figure 6D).

### Expression of TE LINE-1 Is Associated With Male Differentiation and not Apoptosis

We next asked if clonally high ORF1p is associated with germ cell apoptosis, suggesting that an inability to repress TEs could be the basis for germ cell elimination. Recent studies have shown that ORF1p expression is similarly heterogeneous in female oocytes at E15.5 and predictive for DNA damage and eventual oocyte death (Malki et. al., 2014). Surprisingly, we found no significant correlation between high ORF1p and apoptosis in males during this specific apoptotic wave (Fig S5). Our observation of the accumulation rather than elimination of ORF1p-high cells from E12.5 through E15.5 further argues against apoptotic coordination with ORF1p expression. To assess heterogeneity of TEs in relation to transcriptional identities, we aligned our single-cell RNA data to a transposon reference genome (Jin et. al., 2015, Reznik et. al., 2019). We used the clustering information from the initial transcriptional analysis to discover TEs that were differentially expressed among the nine detected subpopulations at E13.5. While few transposons significantly marked any of the subpopulations queried (Figure S6), *L1MdA* uniquely identified MD-germ cells (Figure 6E). *L1MdA* is the youngest and most active LINE-1, which comprises the largest class of TEs representing 17% of the genome. Recent evidence has shown that *L1MdA* follows the same demethylation and expression dynamics as GRRs (Hill et. al., 2018). Since GRR expression reflects a pro-differentiation epigenetic state, the expression of *L1MdA* provides further evidence of the distinct epigenetic states differentiating MD-germ cells from AP-germ cells.

### Apoptosis Eliminates LINE-1 Negative Germ Cells

Increased expression of ORF1p in germ cells from E13.5 to E15.5 occurs concomitantly with male differentiation, concluding with a largely homogeneous ORF1p-positive MD-germ cell population. Such developmental homogeneity is likely achieved through the elimination of developmentally aberrant clones, as we have demonstrated with apoptosis targeting AP-germ cells that improperly maintain *Lefty* expression. To evaluate the contribution of apoptosis to promoting MD-germ cells, we investigated the expression of ORF1p in *Bax* null embryos where inhibition of apoptosis retains aberrant germ cells. While wild-type germ cells almost entirely express ORF1p at E15.5, *Bax* mutants contained a significantly higher number of germ cells that were deficient for ORF1p (Fig 7A, B). These ORF1p-negative germ cells also were organized in clusters, which could represent clonally related cells sharing a differentiation defect preventing appropriate activation of LINE-1 and ORF1p at E15.5.

**Figure 7.**
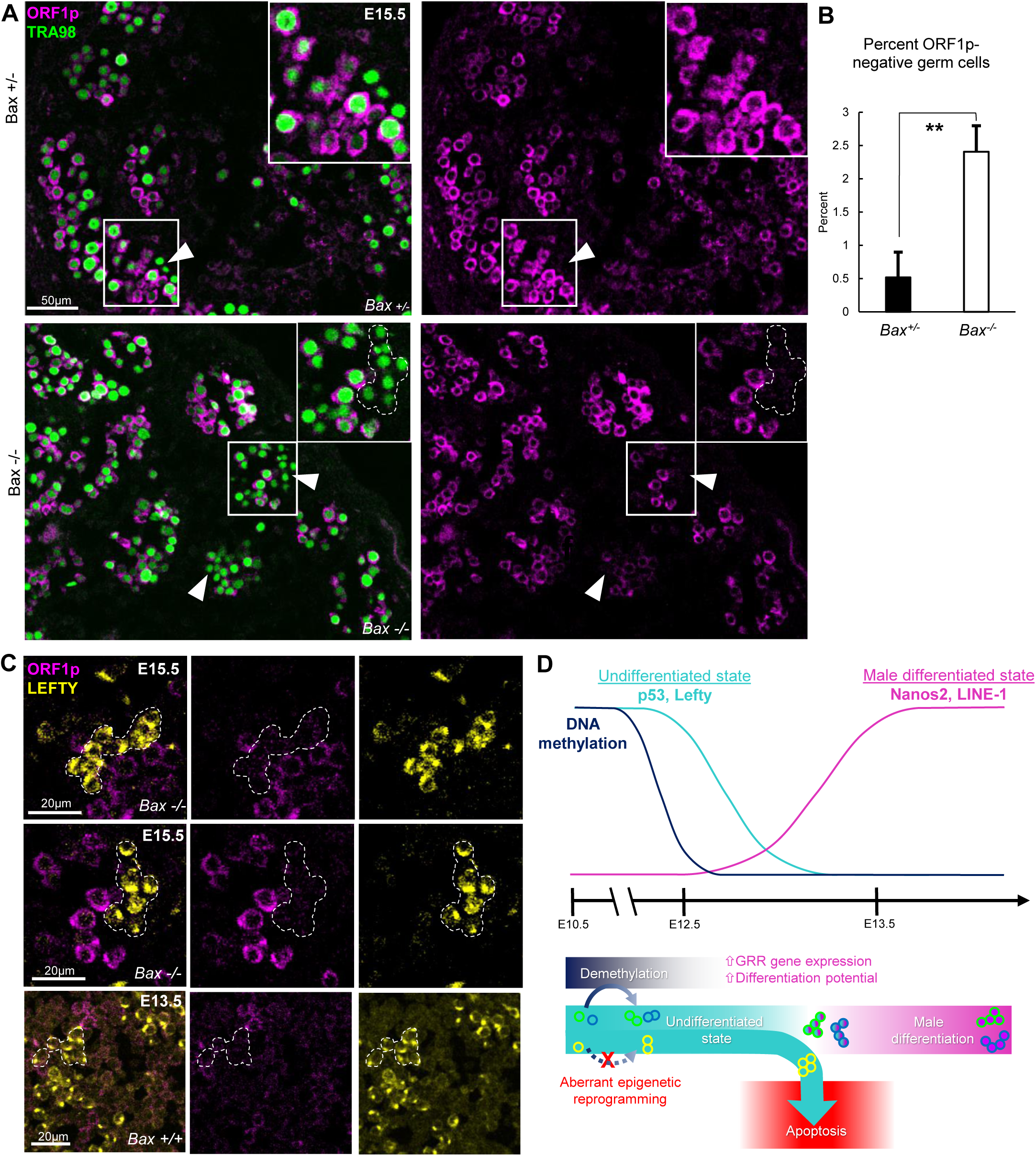
LINE1 is absent in aberrantly differentiated germ cells. **A**, Persistence of ORF1p-negative germ cells in the absence of apoptosis in E15.5 testes. Comparison of *Bax* heterozygote to homozygote knockouts. ORF1p-negative germ cells indicated by arrowheads. Inset: higher-magnification image of ORF1p-negative groups of germ cells with a cluster outlined. **B,** Comparison of percentage of ORF1p-negative germ cells in Bax homozygote knockouts versus heterozygotes at E15.5. N=2 for each genotype. ** = p<0.05. **C,** ORF1p expression is reciprocal to immature differentiation marker LEFTY. LEFTY-positive germ cells in *Bax* mutants at E15.5 are ORF1p negative. Similar reciprocal expression shown at E13.5 in wild-type (bottom). LEFTY-positive clones outlined. **D.** Model for clonal origin of heterogeneous differentiation leading to clonal apoptosis or survival. Top: Expression dynamics of male-differentiation genes following demethylation and epigenetic reprogramming. Demethylation, particularly at GRRs, precedes the transition to a male-differentiated state. Bottom: Model for clonal divergence due to differences in epigenetic reprogramming. Demethylation at GRRs facilitates differentiation and successful survival whereas aberrant epigenetic reprogramming disrupts differentiation-related gene expression and results in these germ cell clones dying.

### LINE-1 is Absent in Aberrantly Differentiated Germ Cells

Considering the similar clustering of LEFTY-positive AP-germ cells that are increased in *Bax*, we hypothesized that the ORF1p-negative germ cells may be the same aberrant germ cell subpopulation. We examined the expression of LEFTY in the ORF1p-negative germ cell clusters and verified that LEFTY-positive cells at E15.5 are indeed ORF1p-negative (Figure 7C). Taken together, these results argue that inverse expression of LEFTY and ORF1p defines two opposite differentiation states that lead to divergent germ cell fates: inappropriately sustained LEFTY expression is associated with aberrant differentiation (AP-germ cells) and apoptosis, while ORF1p expression reflects progression to a male-differentiated (MD-germ cell) state that survives.

## Discussion

Successful reproduction and genetic transmission is dependent on the appropriate sex differentiation of germ cells. Here we show that apoptosis nonrandomly selects among heterogeneous germ cell populations in the fetal mouse testis to promote appropriate male differentiation. We utilize wholemount confocal imaging with multicolor clonal labeling to reveal that apoptotic germ cell clusters are comprised of clonally related cells. Using single-cell RNA sequencing, we identify an apoptosis poised (AP) germ cell state distinguished by elevated expression of *Trp53* and other genes associated with a sexually-undifferentiated state. We also identify a contrasting, apoptosis-resistant state defined by male-differentiation (MD) and expression of *Nanos2*. Together, these two germ cell states represent a dichotomy between apoptosis and male differentiation that defines winners and losers in fetal germ cell development. These insights from our single-cell approach underscore the utility of investigating a singular cell type during a differentiation process to uncover developmental heterogeneity and its impacts on subpopulation outcomes. We also show that expression of epigenetically-regulated genes such as LINE-1 further defines MD-germ cells, suggesting that epigenetic reprogramming differences are a source of the heritable variations underlying clonally heterogeneous behaviors like apoptosis. Together, these findings suggest that the wave of fetal apoptosis effectively enriches the overall quality of the spermatogonial progenitor pool by removing less developmentally competent cells.

The elimination of aberrantly differentiated germ cell clones may be beneficial toward ensuring reproductive success. *Bax* male mutants are infertile and demonstrate meiotic errors, but here we show that defects in GCs are already apparent in the fetal period far earlier than the adult stage when fertility is examined. Using a single-cell RNA-seq approach on a population of germ cells during the fetal apoptotic wave, we are able to discern and characterize an apoptosis-poised population for potentially aberrant gene expression. We find that the AP-germ cell state identified by elevated Nodal signaling and inappropriate expression of immaturity markers such as *Lefty* and *Rhox6/9* is also deficient in male differentiation. By E15.5 when male differentiation is ordinarily completed, AP-germ cells still fail to reach the MD-germ cell male state even when afforded a longer differentiation window in *Bax* mutants, where AP-germ cells are never eliminated. The continued inability to differentiate suggests that the initial defect associated with AP-germ cells is not merely a transient delay. Rather, this defect is robust and maintains cells in an undifferentiated state with unabated Nodal signaling. Nodal is associated with pluripotency in germ cells (Spiller et. al., 2012) and Nodal-high cells have higher tumorigenic potential (Watanabe et. al., 2010). In the germline specifically, high Lefty and Nodal expression is found in carcinoma *in situ* tumor stem cells that produce testicular germ cell tumors (TGCT) (Spiller et. al., 2013). Apoptosis may function to remove germ cells with aberrantly persistent Nodal signaling to prevent the initiation of germ cell tumors upon defective differentiation. While TGCTs were not reported in *Bax* mice, the tumorigenic effects of inhibiting germ cell quality control may be more evident on a TGCT-prone background (Dawson et. al., 2018).

Recent studies demonstrate that epigenetic regulation at a specific set of sex-differentiation genes determines whether a cell can efficiently proceed to a male identity. The promoters at recently identified germline reprogramming-responsive (GRRs) genes must be demethylated to enable expression (Hill et. al., 2018). Mouse as well as human germ cells undergo significant demethylation and epigenetic reprograming between E7.5 and E12.5 prior to (2) sex-differentiation at E12.5., although the synchronicity and homogeneity of reprogramming across all fetal germ cells is unclear. Epigenetic marks are preserved through cell divisions and any variation in GRR demethylation could propagate clonally to produce progeny with distinct differentiation capabilities. Our observation of clustered AP cells with aberrant male differentiation supports this model of clonal epigenetic variation leading to distinctive germ cell fates. We show that AP germ cells also are deficient for several GRR genes, supporting the notion that aberrant epigenetic reprogramming underlies the AP state. The origins of this clonal epigenetic diversity are presently unknown. Combining clonal labeling with bisulfite sequencing can potentially uncover loci-specific epigenetic differences that generate germ cell heterogeneity. The critical period of fetal epigenetic reprogramming also precedes female sex differentiation and subsequent fetal female apoptosis as well (de Felici and Klinger, 2015). Although the causes for female apoptosis are unclear, aberrant epigenetic reprogramming may also generate a female equivalent of the AP subpopulation that is eventually fated for elimination.

Our finding that the retrotransposon LINE1 is uniquely expressed in MD-germ cells further supports an epigenetic basis for heterogeneous success in differentiation. Similar to GRR genes, the expression of LINE1 is upregulated upon demethylation at its promoter, hence the levels and timing of LINE1 can likewise indicate the extent of epigenetic reprogramming to enable male differentiation. We observe the highest LINE1 in MD-germ cells, suggesting that this subpopulation is the most epigenetically poised for efficient male differentiation. Conversely, AP-germ cells defined by persistent *Lefty* expression are deficient for LINE1, as evidenced in the accumulation of LINE1-negative germ cells when elimination of AP-germ cells is blocked in *Bax* mutants. Altogether, these results support a paradigm in which heterogeneity in epigenetic reprogramming at GRR loci produces a diversity of clones with variable capabilities for male differentiation (Figure 7D). The resulting clonal differences can lead to disparate cell fates, with apoptosis and male differentiation as two diametrically opposed outcomes.

The association of LINE-1 TE expression with the MD-germ cell state was surprising considering the enrichment in these cells for piRNA-biogenesis genes such as *Mili* and *Mael*. Mutations in piRNA regulatory machinery such as Miwi2, Mael, Mov10l, and Dnmt3L lead to overexpression of TEs and reproductive failure during later spermatogenesis (Aravin et. al., 2008; Soper et. al., 2008; Zheng et. al., 2010; Bourc’his and Bestor, 2004). Considering these phenotypes, it would be expected that piRNA-high MD-germ cells would exhibit TE suppression. However, our results suggest the opposite: MD-germ cells show elevated levels of LINE-1 TEs as measured by ORF1p and the proportion of LINE-1-positive germ cells increases throughout the apoptotic wave. The decrease in *Trp53* that is associated with MD-germ cells may partially explain this result. As P53 is known to inhibit TEs, (Wylie et. al., 2016), it is possible that incipient LINE-1 expression is observed as germ cells transition to a P53-low MD-germ cell state. Whether the expression of LINE-1 in fetal germ cell development plays a functional role remains unclear. While LINE-1 is ultimately repressed by piRNAs in adult germ cells, its early expression in the fetal period could be beneficial to priming the piRNA system. A mechanism of piRNA biogenesis in the fetal testis involves an amplification loop termed the ping-pong cycle in which sense transcripts from active transposons guide piRNA amplification and direct methylation at TEs for more durable repression (Aravin et. al., 2008). The initially high expression of LINE-1 associated with newly differentiated MD-germ cells suggests that this population would be first to initiate ping-pong biogenesis and subsequently maintain lower levels of LINE1 throughout adulthood. Hence, the survival advantage associated with MD-germ cells may enrich for germ cells with more robust piRNA production and improved long-term suppression of TEs. In human germ cells, elevated TE expression has also been observed in an advanced male subpopulation, further confirming that heterogeneity among germ cells can manifest as population-specific distinctive TE expression (Reznik et. al., 2019). The advanced human fetal germ cells that also express relatively higher piRNA genes demonstrate later repression of TEs, which is consistent with active PIWI-piRNA silencing of TEs following ping-pong amplification. Future investigations that follow the earliest differentiating germ cells such as MD-germ cells beyond the fetal period can determine how primacy in male differentiation improves TE regulation and genomic integrity.

## Acknowledgements

Rainbow mice were gift from I.L. Weissman. Tex14 mice were a gift from M. Matzuk. ORF1p antibody was a gift from A. Wilkin. We are grateful for feedback and comments from colleagues J. Sneddon, M. Conti, T. Nystul, A.W. De Tomaso, L. Byrnes, D. Wong and all members of the Laird lab including R.G. Jaszczak, S. Cincotta, B. Reznik, S. Peng, and assistance from Jennifer Daza, Gul Bikem Soygur, Grace Zhang, and Ben Dreskin. This study was funded by NSF Predoctoral Fellowship to D.H.N and NIH 1DP2OD007420, R01 GM122902, and Cancer Research Coordinating Committee grant to D.J.L.

## Author Contributions

D.N. designed and performed the experiments and wrote the manuscript. D.L. designed and oversaw all experiments and wrote the manuscript.

## Declaration of Interests

The authors declare no competing interests.

## Methods

### Mice

For WT embryo collection, CD1 females were mated to Oct4-ΔPE-GFP^Szabo^ males (MGI: 4835542). For clonal labeling, *R26R-Confetti*^*Snippert*^ (MGI:104735) and *R26R-Rainbow*^*Rinkevich*^ mice (gift from I. Weissman, Stanford University) were outcrossed onto CD1 to generate mixed background homozygous females and then crossed to heterozygous *Pou5f1-cre/Esr*^Greder^ (MGI:5049897) males for Tamoxifen-inducible germ-cell specific labeling after e8.5. *Tex14*^tm1Zuk^ (*Tex14*^-/-^) mice were a gift from M. Matzuk (Baylor College of Medicine).

### Wholemount and Section Imaging

Tissues were fixed in 4% PFA for 2h, washed with PBS, and blocked with 2% BSA, 0.1% Triton X-100 in PBS for 3 hours. Primary antibodies incubation was performed in 0.2% BSA, 0.1% Triton X-100 in PBS for 2 or more days at 4°C, followed by washing with 0.1% Triton X-100 in PBS. Primary antibodies used were Tra98, Abcam (ab82527), 1:200; cleaved-PARP Alexa 647-conjugated, BD Biosciences(F21-852), 1:20; cleaved-PARP, Cell Signaling (9544), 1:100; P53, Cell Signaling (2425S), 1:100

Secondary antibody incubation was performed in 0.2% BSA, 0.2% BSA, 0.1% Triton X-100 in PBS. Tissues were washed with PBS and dehydrated through a 25%, 50%, 75%, 100%, 100% methanol series. Tissues were cleared with a 2:1 benzyl benzoate:benzyl alcohol (BABB) solution and imaged in BABB with a *10x*/0.4 *dry* HCX PL APO CS objective on a Leica SP8 upright confocal microscope.

For section immunofluorescence, tissues were fixed in 4% PFA for 2h, washed with PBS, and dehydrated overnight in 30% sucrose at 4C. Tissues were embedded in OCT and flash-frozen and stored at −80C. Thick cryosections were cut at 25um and 50um; otherwise, sections were cut at 8um thickness and affixed to Superfrost Plus slides (Fisher Scientific). Sections were washed with PBS and incubated overnight at 4C with primary antibody in 5% donkey serum, 0.5% Triton X-100. Sections were washed with PBS and incubated with secondary antibody for 1h at room temperature. Slides were mounted with Vectashield and imaged on a SP5 Leica confocal microscope. For antibodies requiring antigen retrieval, sections were immersed in 10mM sodium citrate and heated until boiling. Sections were washed with PBS and stained with primary antibody as described.

### RNA in situ hybridization

Testes were prepared for section immunofluorescence and sectioned at 5µm. RNA was detected using the RNAscope Multiplex Fluorescent kit v2 with probes against mouse *Mm-Tp53-C2* and *Mm-Nanos2-C1*. Tissue sections were pretreated with RNAscope Protease III for 6 minutes at 37°C and incubated in hydrogen peroxide for 10 minutes at 37°C. Probes were incubated for 2 hours followed by the standard Multiplex Fluorescent v2 assay. Probes were detected by Opal dyes 570/650 (Perkin Elmer). For subsequent antibody-based immunofluorescence, sections were prepared using the described section immunofluorescence protocol.

### Spatial statistical analysis

Wholemount tissues were stained for makers of apoptosis, GCs, and nuclei. Objects were identified using the Find Objects function in Volocity (PerkinElmer Improvision) to determine the three-dimensional coordinates of each object centroid. Spatial analysis of clustering was based on the Ripley K function using the RipleyGUI (12) platform in Matlab. K-function scores were calculated to evaluate deviation (K(t) – *E*[K(t)]) from an expected random distribution, CSR, which was simulated independently 100 times for each spatial distribution analyzed. The relative degree of clustering for apoptotic GCs versus all GC distributions, or across time points, was tested with the between-treatments sum of squares (BTSS) and compared to a 95% confidence interval for the bootstrapped BTSS value of the null hypothesis.

### Clonal analysis

*R26R-Confetti* and *R26R-Rainbow* female mice were mated with *Pou5f1-cre/Esr* males and intraperitoneally injected with Tamoxifen (Sigma, 20mg/ml dissolved in sunflower seed oil) at E10.5. Tamoxifen dosage was scaled to the pregnant female’s weight and adjusted to produce distinguishable colored populations (1.25mg and 2.5mg/40g female for *Confetti* and *Rainbow*, respectively). Clonally labeled gonads were dissected and fixed for wholemount staining or section immunofluorescence as described.

To clear tissues and preserve endogenous fluorescence for wholemount imaging, tissues were washed with PBS following secondary antibody incubation and placed in Scale CUBIC Reagent 1^Susaki^ overnight. Cleared tissues were imaged in Scale CUBIC Reagent 1 on a white-light Leica SP8 confocal microscope. Excitation for CFP was with a 458nm laser line; GFP and YFP, 514nm white-light; RFP, 561 white-light. Fluorescence was collected for CFP between 465-495nm, airy 1.5; GFP and YFP, 521-555nm; RFP, 565-590nm.

Clonal populations were analyzed using the Cell module on Imaris to identify individual cells of a clone and quantify clone size. Clones were detected by CFP, YFP, or RFP intensity with a threshold set at 2 standard deviations below the median intensity value. Intensity was measured over the cell body with a 1.2um background filter and a 5um minimum cell diameter. Individual cells were separated using a 7um estimated cell diameter. For clonal identification, similarly colored cells within the dispersion distance of 50um were considered to be part of the same clone.

### Single cell RNA seq

For E12.5 and E13.5 male GC collection, WT testes from E12.5 and E13.5 timed matings were collected together. Testes were dissected in cold PBS and non-gonadal tissue removed. Testes were digested in 0.25% trypsin/EDTA at 37C for 20 minutes with trituration every 10 minutes, followed by the addition of 1mg/ml DNAse and further digestion for 10 minutes. An equal volume of fetal bovine serum was added to halt digestion and the digest was strained through a single-cell filter. Dead cells were labeled with Sytox Blue and live GCs were obtained by sorting on GFP^+^, Sytox^-^ into 0.04% BSA. 2,517 and 2,606 cells were recovered for E12.5 and E13.5 timepoints, respectively. Cells were processed for 10X sequencing by the UCSF Institute for Human Genetics. Cell by gene matrices were obtained by performing CellRanger analysis on 10x reads.

Single cell expression data was analyzed using Seurat to identify differentially expressed genes and perform principal component analysis (35). Statistically significant principal components (n=17, p<0.05) were used to cluster cells in an unsupervised manner. Differentially expressed genes by cluster (GC state) were identified by receiver operating characteristic (ROC) test, bimodal test, and Wilcoxon rank sum test. Significance cutoffs were AUC>0.6, ROC test and p<0.05 for bimodal and Wilcoxon rank sum test. An AUC score>0.6 was used to identify markers that were significant, positive classifiers of a cluster. Fold changes and normalized gene expression was expressed in natural log space as a default output of Seurat. For fold change calculations pertaining to cluster markers, the mean expression across all cells of one cluster was compared to the mean expression of all other cells in non-log space, and then the fold difference was expressed in natural log space.

Gene ontology analysis was performed using biomarker lists for each clustered population identified by Seurat using GSEA molecular signature database analysis. Biomarkers were analyzed for statistical overrepresentation in biological process categories and semantically sorted using ReviGO.

For pseudotime analysis, E13.5 GC expression data was analyzed using Monocle (36). We used the unsupervised dpFeature procedure to identify the top 1000 highly variable genes among clusters for constructing the differentiation trajectory. Cells were clustered with a rho of 15 and delta of 8.

### Supplemental Information

scRNA-seq data are available at https://www.ncbi.nlm.nih.gov/geo/query/acc.cgi?acc=GSE119045

### Supplemental movie 1

#### Wholemount visualization of apoptotic clusters of GCs at E13.5

Whole E13.5 testis with cell nuclei identified by Hoechst and GCs by Tra98. Apoptosis is identified by cleaved-PARP and rotated views highlight the spatial distribution of apoptotic GCs. Magnified views show a representative apoptotic GC cluster.

### Supplemental movie 2

#### GC proliferation is clonal

E13.5 GCs are clonally labeled using *Rainbow* and cells in late G2/M are marked by phospho-histone H3. Magnified view shows two separate groups of phospho-histone H3^+^ cells that are clonal (mOrange clone and a GFP^-^ clone).

---

**Supplemental Table 1.** Transcriptional markers that distinguish each single germ cell state compared to all others were identified among differentially expressed genes in an E13.5 germ cell population by the Wilcoxon rank sum test-based FindMarkers operation in Seurat.

**Figure S1.**
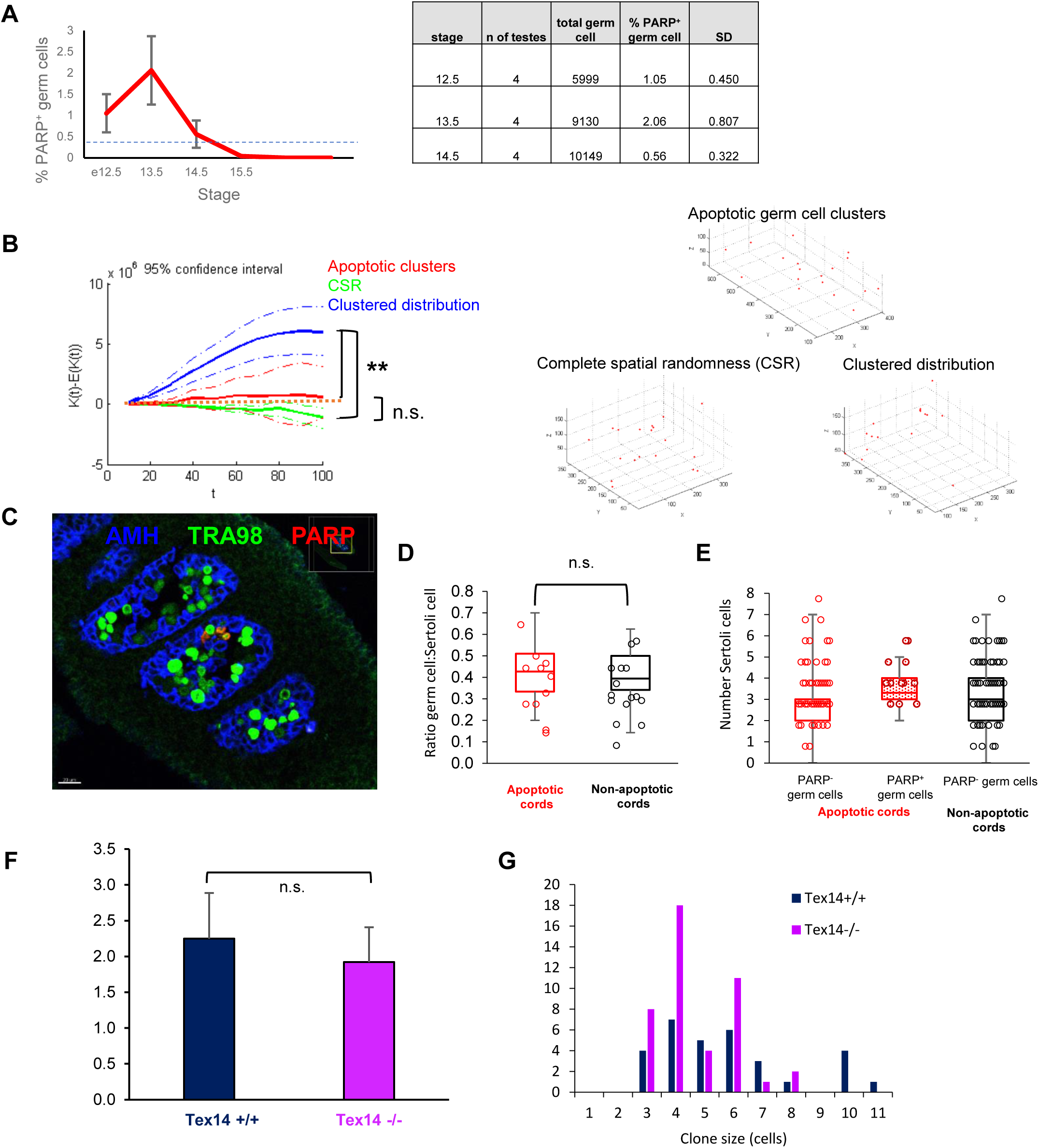
Related to Figure 1. **A,** Apoptotic cleaved PARP^+^ germ cells quantified in wholemount testes during apoptotic wave. Dashed line denotes reported average apoptotic index in developing fetal tissues (REF 12) **B,** Distribution of apoptotic clusters measured for spatial bias (clustering). The 3D coordinates of each cluster’s centroid was plotted for Ripley K-factor analysis. Apoptotic germ cell cluster distribution (left) was compared against simulated distributions for an equivalent number of points under simulated non-clustered (complete spatial randomness, CSR) and clustered distributions. Representative graphs for each example distribution (right) depict deviation from spatial randomness. Apoptotic clustering comparison performed for n=5 e13.5 testes versus 1000 simulations of CSR and clustered distributions for an equivalent number of clusters. **; p<0.001. **C,** Section of e13.5 testis stained for Sertoli cells (AMH), germ cells (TRA98), and apoptosis (cleaved-PARP). Apoptotic cords are defined as cords containing any apoptotic germ cells (red outline) in contrast to non-apoptotic cords (white outline). **D,** Ratio of total germ cells to Sertoli cells in both apoptotic and non-apoptotic cords sections, p-value: 0.584. **E,** Average numbers of Sertoli cells in direct contact with each germ cell in both apoptotic and non-apoptotic cords. Sertoli cell contacts were further distinguished between apoptotic (PARP+) and non-apoptotic (PARP-) germ cells. Apoptotic cords are plotted in red and cords without apoptosis are plotted in black plot. For all 3, p>0.25. **F**, Quantification of apoptotic germ cells in E13.5*Tex14* mutants. **G,** Apoptotic cluster size by number of PARP^+^ germ cells in Tex14+/+ and Tex14^-/-^E13.5 testes

**Figure S2.**
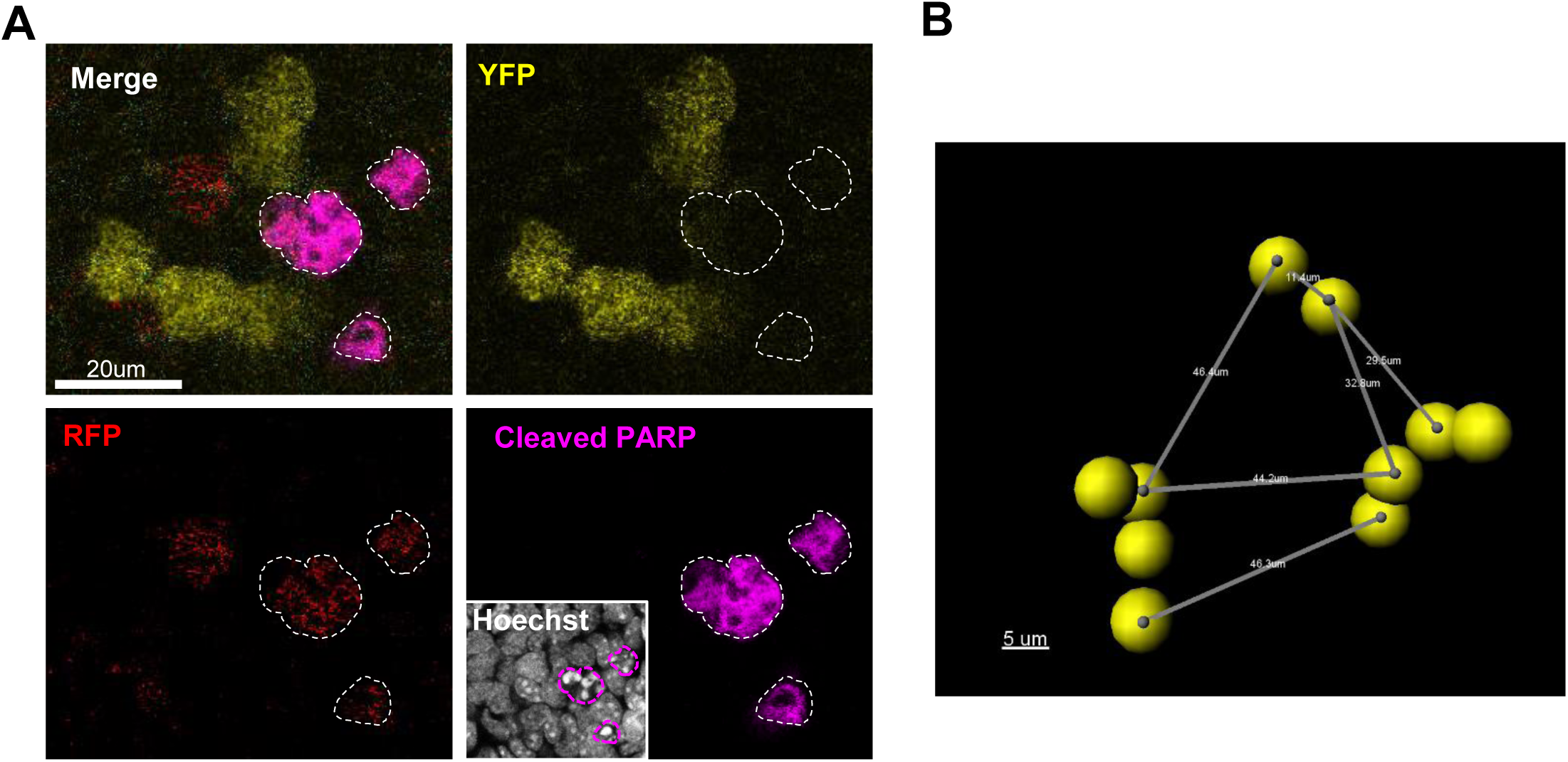
Related to Figure 2. **A,** Detection of clonal apoptosis in *Confetti* labeled E13.5 germ cells. Apoptotic germ cells identified by cleaved PARP staining (white outlines). **B,** Clonality of apoptosis was determined by identifying inducibly-labeled (*Rainbow* or *Confetti*-positive) apoptotic germ cells and recording the number and clonal label of nearby apoptotic germ cells. To define the search radius for considering two similarly labeled apoptotic germ cells to be clonal, we first defined the maximum distance for cells of a clone to disperse from each other. We determined this distance in wholemount by generating isolated germ cell clones in e13.5 *Pou5f1-CreER x Rainbow/Confetti* testes that were induced with tamoxifen at e10.5 (inset). At a low (0.5mg) dose of tamoxifen, clonal labeling was sufficiently rare to generate distinct single clones. Germ cells are immotile after colonizing the gonadal ridge at e10.5 (#32) so clonally related cells are confined to a local boundary. Cells of a single clone can fragment into multiple cysts and has been described in (#14). The maximal dispersion of cells of a single clone specifies this boundary and the volume in which we consider clonal outcomes such as apoptosis among similarly colored cells. The maximal dispersion was considered to be the distance to a nearest neighbor cell of a different fragment. In the above example, 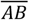 represents the nearest-neighbor relationship for cells A and B but these cells are proximal and are considered as a single fragment. The nearest-neighbor fragment to fragment 1 contains cells C and D but cell C is the nearest-neighbor cell of fragment 2 to fragment 1. Hence, 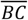 defines the dispersion between these two fragments. Likewise, 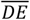 represents the dispersion between fragments 2 and 3 and 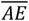 is the dispersion between fragments 1 and 3. The maximal dispersion is therefore 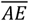, 46.4µm. The mean maximal dispersion among cells of a clone was found to be less than 50 µm, which we established as the range in which we considered similarly colored cells to be clonally related fragments. This distance was corroborated with measurements we obtained from secondary analysis of clonal fragmentation in similarly aged testes (REF Lei) To identify unique apoptotic clusters in wholemount, a group of n>2 PARP^+^ germ cells were considered a cluster if the measured distance between the germ cells was below 50 µm. PARP^+^ germ cells beyond 50 µm from each other were considered to belong to a separate cluster. The median diameter for apoptotic clusters was obtained by measuring the largest distance spanned by two cells of an apoptotic cluster.

**Figure S3.**
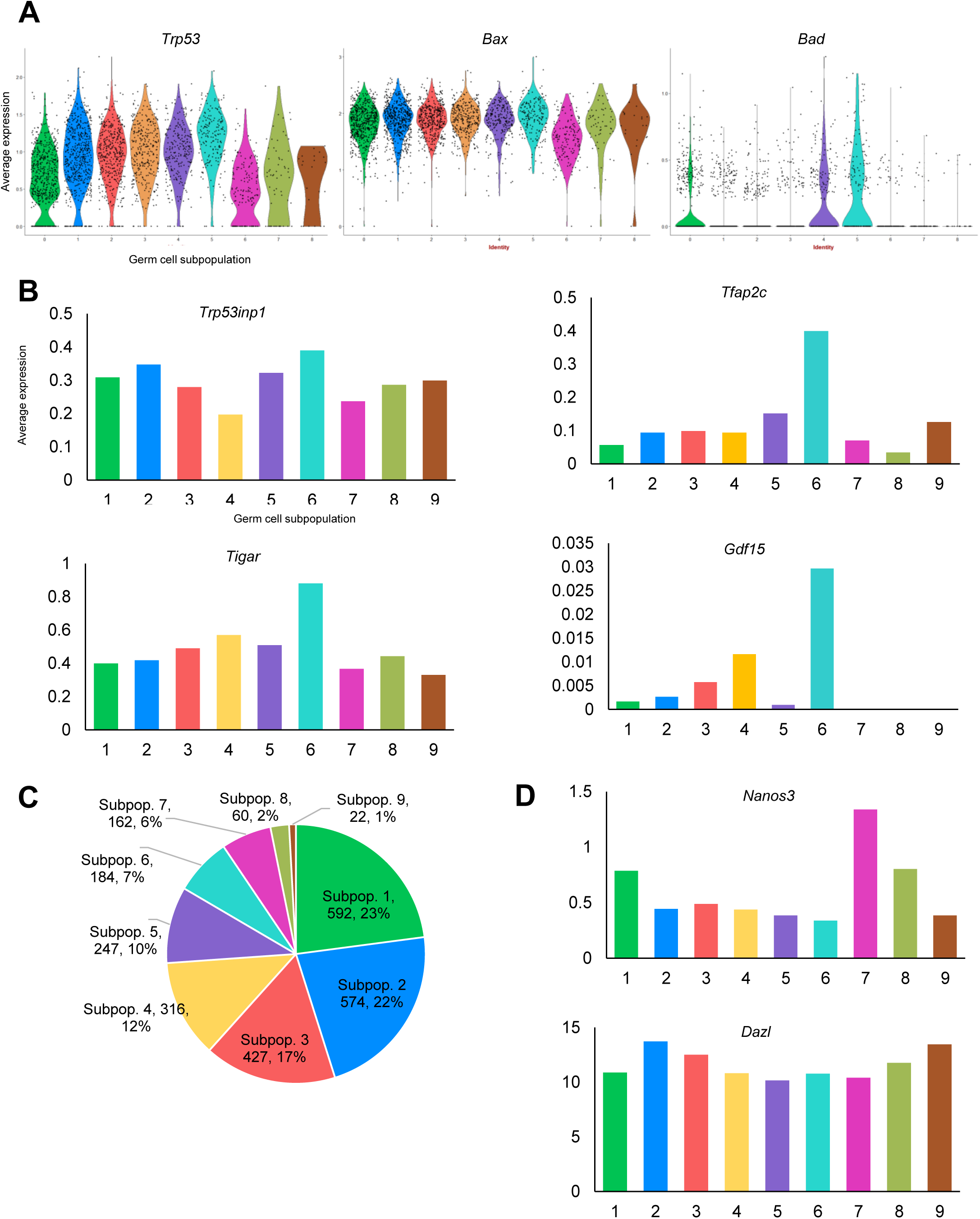
Related to Figure 3. **A,** Violin plots for average expression of AP-germ cell markers by subpopulation. Individual cells are represented as single dots. **B,** Expression of p53-activated genes by germ cell subpopulation. **C,** Subpopulation size in cell number and as a percentage of total germ cells analyzed for single-cell RNA sequencing at E13.5‥ **D,** *Nanos3* expression by germ cell subpopulation. **E,** *Dazl* expression by germ cell population

**Figure S4.**
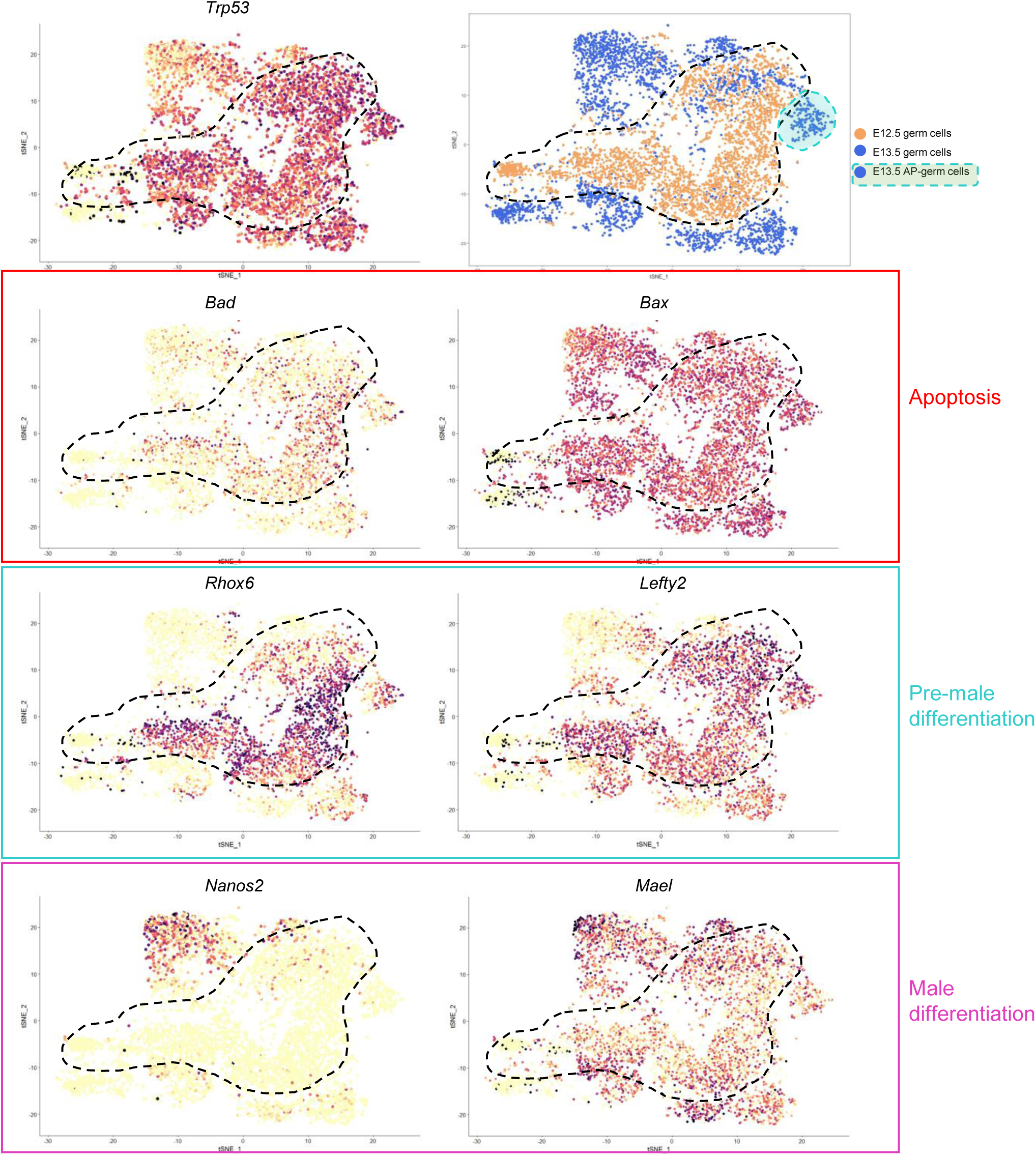
Related to Figure 4. tSNE plots showing expression of genes *(Bad, Bax*), male differentiation genes *(Nanos2, Mael*), and apoptotic genes associated with a pre-male differentiation population *(Rhox6, Lefty2*) in combined e12.5 and e13.5 germ cells. Black dashed line indicates approximate boundary of E12.5 population.

**Figure S5.**
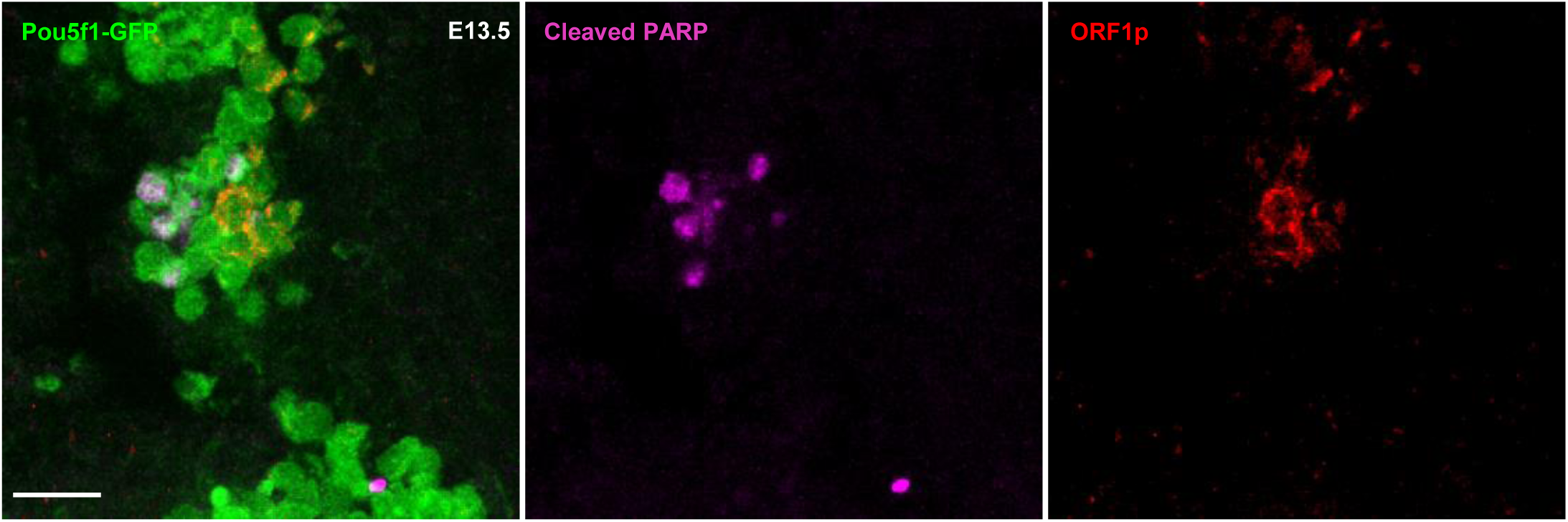
Related to Figure 6. ORF1p and cleaved PARP staining in E13.5 wild type sections

**Figure S6.**
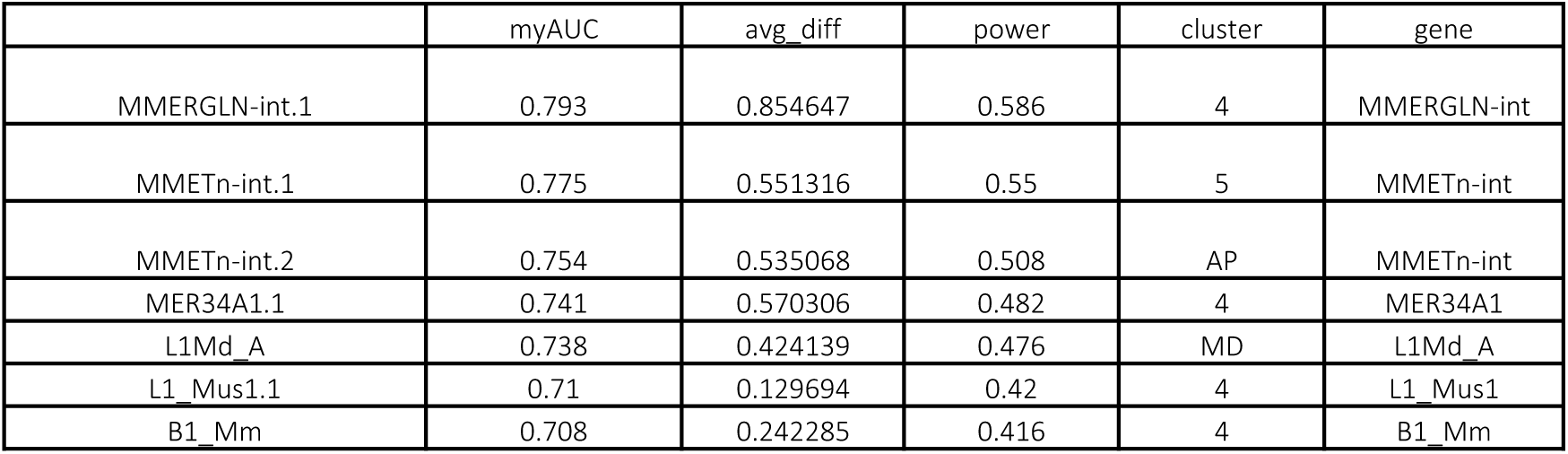
Related to Figure 6. Differentially expressed repeat elements for E13.5 germ cell clusters that were identified by single-cell RNA transcriptional profiling. Markers were determined by ROC test.

